# Single-dose intranasal vaccination elicits systemic and mucosal immunity against SARS-CoV-2

**DOI:** 10.1101/2020.07.23.212357

**Authors:** Xingyue An, Melisa Martinez-Paniagua, Ali Rezvan, Mohsen Fathi, Shailbala Singh, Sujit Biswas, Melissa Pourpak, Cassian Yee, Xinli Liu, Navin Varadarajan

## Abstract

A safe and durable vaccine is urgently needed to tackle the COVID19 pandemic that has infected >15 million people and caused >620,000 deaths worldwide. As with other respiratory pathogens, the nasal compartment is the first barrier that needs to be breached by the SARS-CoV-2 virus before dissemination to the lung. Despite progress at remarkable speed, current intramuscular vaccines are designed to elicit systemic immunity without conferring mucosal immunity. We report the development of an intranasal subunit vaccine that contains the trimeric or monomeric spike protein and liposomal STING agonist as adjuvant. This vaccine induces systemic neutralizing antibodies, mucosal IgA responses in the lung and nasal compartments, and T-cell responses in the lung of mice. Single-cell RNA-sequencing confirmed the concomitant activation of T and B cell responses in a germinal center-like manner within the nasal-associated lymphoid tissues (NALT), confirming its role as an inductive site that can lead to long-lasting immunity. The ability to elicit immunity in the respiratory tract has can prevent the initial establishment of infection in individuals and prevent disease transmission across humans.

## Introduction

The COVID19 pandemic has made the development of an efficacious and safe vaccine an urgent priority. Rapid progress in sequencing, protein structure determination, and epitope mapping with cross-reactive antibodies has illustrated that the SARS-CoV-2 spike protein (S protein) binds to human angiotensin-converting enzyme 2 (ACE2)^1,2^. This, in turn, has made the S protein and the receptor-binding domain (RBD) of the S protein prime candidates for vaccine design to elicit neutralizing antibodies. Indeed, DNA/RNA based vaccine candidates encoded for the S protein have advanced rapidly and are in clinical trials^3,4^.

Despite this progress, there is considerable uncertainty about the duration of protective immunity elicited by the current vaccine candidates. While neutralizing antibodies are the desired immunological target of the current vaccines, and maybe necessary; it is unclear if serum neutralizing antibodies will be sufficient for sterilizing immunity. Studies have shown that the antibody protection in COVID-19 convalescent patients can be short-lived and that patients can become seronegative in as little as four weeks after exposure^5-7^. There are also reports of the lack of high levels of neutralizing activity; and in some cohorts, even a lack of detectable neutralizing antibodies in patient convalescent sera^2,8^.

Adaptive immunity mediated by T cells can complement humoral immunity or can inhibit viral replication independent of humoral immunity. Not surprisingly, there are emerging reports of patients with COVID-19 like symptoms who have detectable T-cell responses without seroconversion^9^. This observation complements other studies that support the existence of robust T cell responses in convalescent patients. Grifoni *et al*. demonstrated that 100 % of non-hospitalized convalescent patients showed antigen-specific CD4^+^ T cells, and 70 % of patients showed antigen-specific CD8^+^ T-cell responses^10^. Weiskopf *et al*. detected strong Th1 type responses directed towards the S protein in COVID-19 patients admitted to the ICU due to moderate to severe acute respiratory distress syndrome (ARDS), ^11^. Similarly, Braun *et al*. have reported CD4^+^ T-cell responses targeting the S protein in 83% of COVID19 patients and a third of healthy patients, presumably due to cross-reactivity to other viruses^12^. Taken together, these data highlight that vaccines that target both humoral and cellular responses can deliver lasting protective immunity.

The nose and the upper respiratory tract are the primary routes of entry for inhalation pathogens like SARS-CoV-2. Not surprisingly, the nasal compartment showed particular susceptibility to SARS-CoV-2 infection and can serve as the initial reservoir for subsequent seeding of the virus to the lung^13^. Consequently, pre-existing immunity within the respiratory tract is highly desirable to prevent pathogen invasion. Despite the well-recognized role of mucosal immunity, most vaccines are designed to elicit circulating humoral immunity without necessarily enabling mucosal immunity. Mucosal vaccination can stimulate both systemic and mucosal immunity and has the advantage of being a non-invasive procedure suitable for immunization of large populations. However, mucosal vaccination is hampered by the lack of efficient delivery of the antigen and the need for appropriate adjuvants that can stimulate a robust immune response without toxicity. The identification of the cyclic GMP-AMP Synthase (cGAS) and the stimulator of interferon genes (STING) pathway has enabled the identification and development of STING agonists (STINGa)^14^. STINGa function as novel immunostimulatory adjuvants for mucosal vaccines against respiratory pathogens, including influenza and anthrax, in mice^15-17^.

In this report, we encapsulated the STINGa, cyclic guanosine monophosphate–adenosine monophosphate (2′3′-cGAMP or cGAMP) in liposomes^17^. We used it as the adjuvant for intranasal vaccination with the trimeric or monomeric versions of the S protein. Our results show that the candidate vaccine formulation is safe and elicits systemic immunity (neutralizing antibodies), cellular immunity (spleen and lung), and mucosal immunity (IgA in the nasal compartment and lung, and IgA secreting cells in the spleen). To the best of our knowledge, we report the first COVID-19 vaccine candidate that elicits mucosal immunity and supports further translational studies as an intranasal non-viral candidate that can induce systemic immunity and confers immunity at the primary site of viral entry.

## Results

### An efficient and stable liposomal adjuvant containing STINGa

To facilitate efficient priming of the immune system within the respiratory compartment, we encapsulated the STINGa within negatively charged liposomes (Figure 1A)^17^. The adjuvant encapsulated liposomes were prepared using a passive drug loading method by hydrating the lipid dry films in buffered solutions containing cGAMP as the STINGa. We removed the free STINGa via ultrafiltration, and the encapsulation efficiency of STINGa was determined to be 35 % by calibration against a standard curve (Figure S1A). Dynamic light scattering analysis showed that the mean particle diameter by intensity of STINGa-liposomes was 81 nm, with a polydispersity index of 0.24, while the the size of blank liposomes was 110 nm (Figure 1B-C). The mean zeta potential of liposomes was negative both with (−35 mV) and without (−68 mV) encapsulated STINGa (Figure 1D-E). We tested the stability of the STINGa-liposomes and showed that they were stable for up to two months at 4 °C, as evidenced by the conservation of particle sizes and the absence of aggregates (Figure S1B). The surface charge of liposomes was also unaltered (−40 mV) after this period (Figure S1B). Collectively, these results established that the negatively charged liposomes had efficiently encapsulated the STINGa and had good stability.

**Figure 1.**
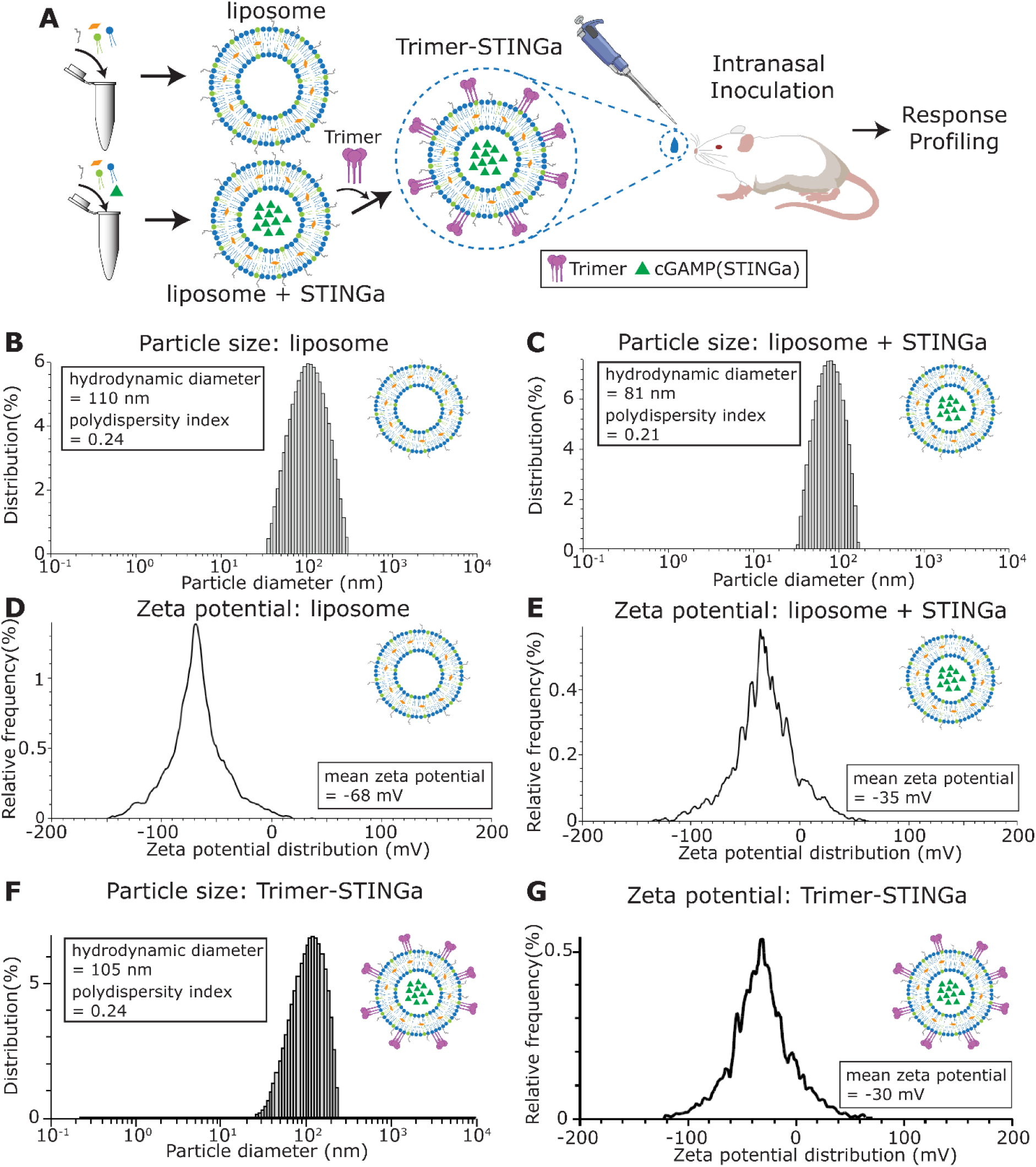
Preparation and characterization of liposome encapsulated STINGa. A. Overall schematic of design of adjuvant and intranasal administration of vaccine. B, C, and F. Distribution of liposomal particle sizes measured by dynamic light scattering (DLS). D, E, and G. Zeta potential of the liposomes measured by electrophoretic light scattering (ELS).

We prepared the vaccine by gently mixing the trimeric S protein (discussed below) with the STINGa-liposomes suspensions at room temperature to allow the adsorption of the protein on the liposomes. The adsorbed Trimer-STINGa-liposomes displayed mean particle diameter of 105 nm and mean zeta potential of -29 mV (Figure 1F-G), with a polydispersity index of 0.24, slightly bigger and less negative than the STINGa-liposomes (81 nm, -35mV). The results suggested that formulated protein-STINGa-liposomes vaccine exist in a nanoparticulate colloidal form.

### Neutralizing antibodies, T-cell responses, and IgA elicited upon vaccination with Trimer-STINGa

We used the recombinant trimeric extracellular domain of the S protein containing mutations to the Furin cleavage site as the immunogen (Figure 2A). As expected by extensive glycosylation of the S protein, SDS-PAGE under reducing conditions confirmed that the protein migrated between 180-250kDa (Figure 2B). Although previous studies have performed extensive characterization of the lack of toxicity of the adjuvant formulation, we wanted to confirm that the adjuvant does not cause morbidity, weight loss, or other hyper-inflammatory symptoms^17^. Accordingly, we performed an initial pilot experiment with a five BALB/c mice that received a single intranasal dose of the adjuvant without protein and observed no weight loss or gross abnormalities over 14 days (Figure S2A and S2B). We next immunized two groups of mice by intranasal administration with either a combination of the protein and adjuvant (Trimer-STINGa) or the protein by itself (control). None of the animals showed any clinical symptoms, including loss of weight (Figure S2C). Seven days (d7) after immunization, 100 % of the mice that received the Trimer-STINGa seroconverted and robust anti-S IgG levels with mean dilution titers of 1:1,040 were detected (Figure 2C). By day 15 (d15), the serum concentration of the anti-S IgG antibodies increased, and mean dilution titers of 1:4,400 were detected (Figure 2C). Anti-S IgG was also detected in the bronchoalveolar lavage fluid (BALF) of all three mice tested (Figure 2D). We confirmed that the serum anti-S antibodies were neutralizing with a mean 50% inhibitory dose (ID50) of 1:414 as measured by a GFP-reporter based pseudovirus neutralization assay (SARS-CoV-2, Wuhan-Hu-1 pseudotype) [Figure 2E].

**Figure 2.**
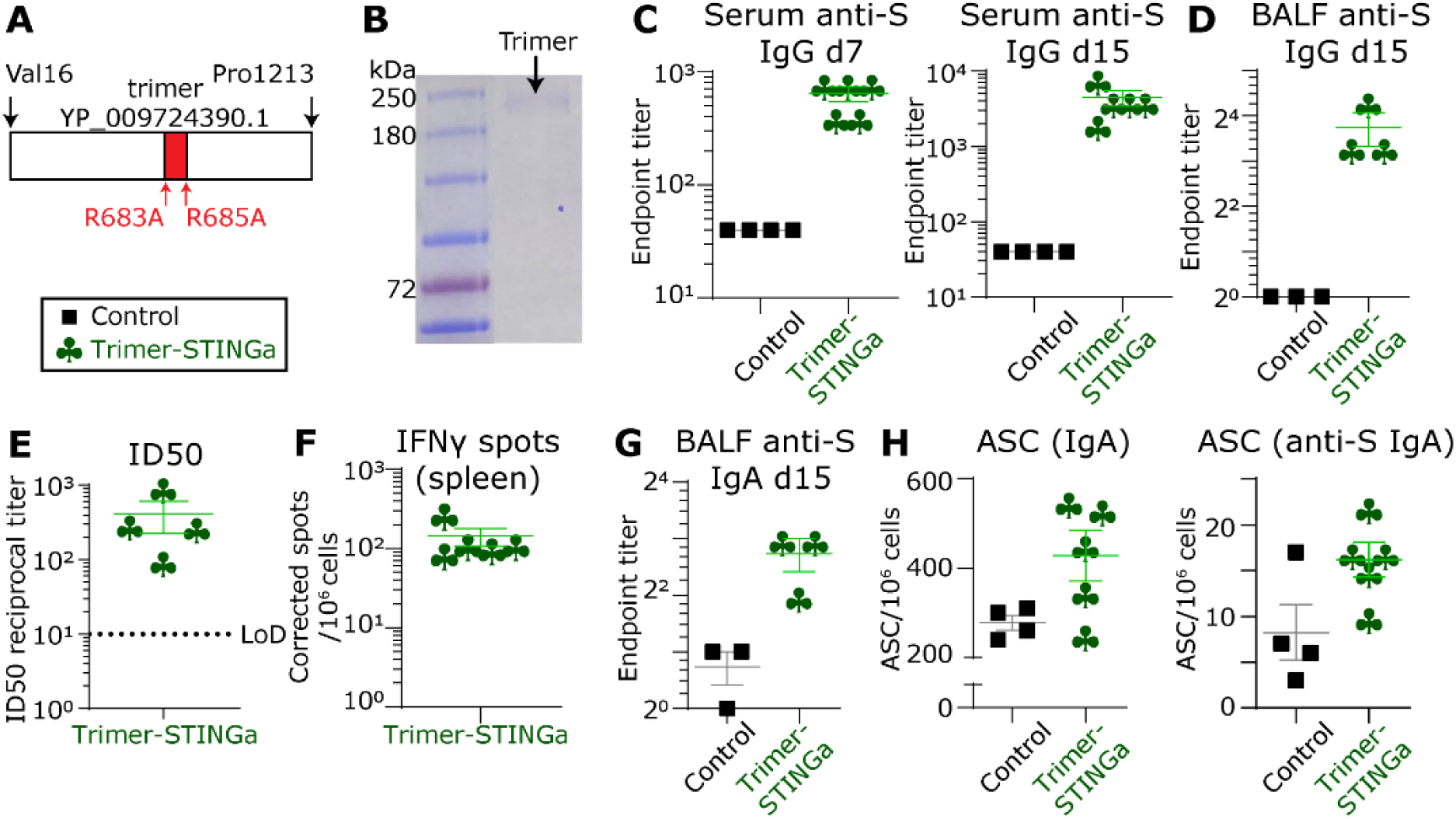
Systemic and mucosal responses elicited upon vaccination with Trimer-STINGa. A. Schematic of trimeric protein used for immunization. B. Denaturing SDS-PAGE gel of the purified trimer protein. C. Humoral immune responses in the serum were evaluated using S-protein based IgG ELISA at day 7 and day 15 after immunization. D. The humoral immune responses in the BALF evaluated using S-protein based IgG ELISA at day 15. E. The ID50 of the serum antibody responses were measured using a pseudovirus neutralization assay F. Cellular immune responses in the spleen were assessed using IFNγ ELISPOT assays. G. IgA levels in the BALF were determined using ELISA. H. Antibody secreting cells (ASCs) secreting IgA and S-protein specific IgA in the spleen were detected using ELISPOT assays.For (C-H) the bar represents the mean, and the error bars represent the standard error. LoD represents limit of detection of assay.

Emerging data support a role for T-cell responses to contribute to protection independent of antibody responses. We evaluated T-cell responses in the immunized mice using a pool of 15mers that target highly conserved regions of the S-protein (Figure S3)^18,19^. At d15, all five animals immunized with the Trimer-STINGa showed robust splenic T cell responses with a mean of 144 IFNγ spots/10^6^ cells (Figure 2F). Collectively these results show that a single intranasal administration using the Trimer-STINGa elicited robust serum neutralizing antibodies and T-cell responses.

IgA mediated protection is an essential component of mucosal immunity for respiratory pathogens. The Trimer-STINGa improved BALF IgA titers at d15 compared to the control group (Figure 2G). We also evaluated the IgA responses in the antibody-secreting cells (ASCs) in the spleen at d15 by ELISPOT assays. The mice immunized with Trimer-STINGa showed an increase in the number of total IgA secreting and S-specific IgA secreting ASCs compared to the control group (Figure 2H). Taken together, these results illustrate that intranasal vaccination elicited IgA responses that are an essential component of mucosal immunity.

### Single-cell RNA-sequencing (scRNA-seq) confirms the nasal-associated lymphoid tissue (NALT) as an inductive site upon vaccination

To investigate if intranasal vaccination can support local inductive responses in the nasal passage, we harvested the NALT from the immunized animals at the time of euthanasia, converted them into single-cell suspensions, and performed scRNA-seq (Figure 3A and S4). After filtering, we obtained a total of 1,398 scRNA-seq profiles. By utilizing uniform manifold approximation and projection (UMAP), we identified the myeloid; NK and T; and B cell subpopulations using established lineage markers (Figure 3B and S5). >95 % of the scRNA-seq in both control and Trimer-STINGa groups corresponded to T and B cells, and we performed detailed analyses on these immune cells.

**Figure 3.**
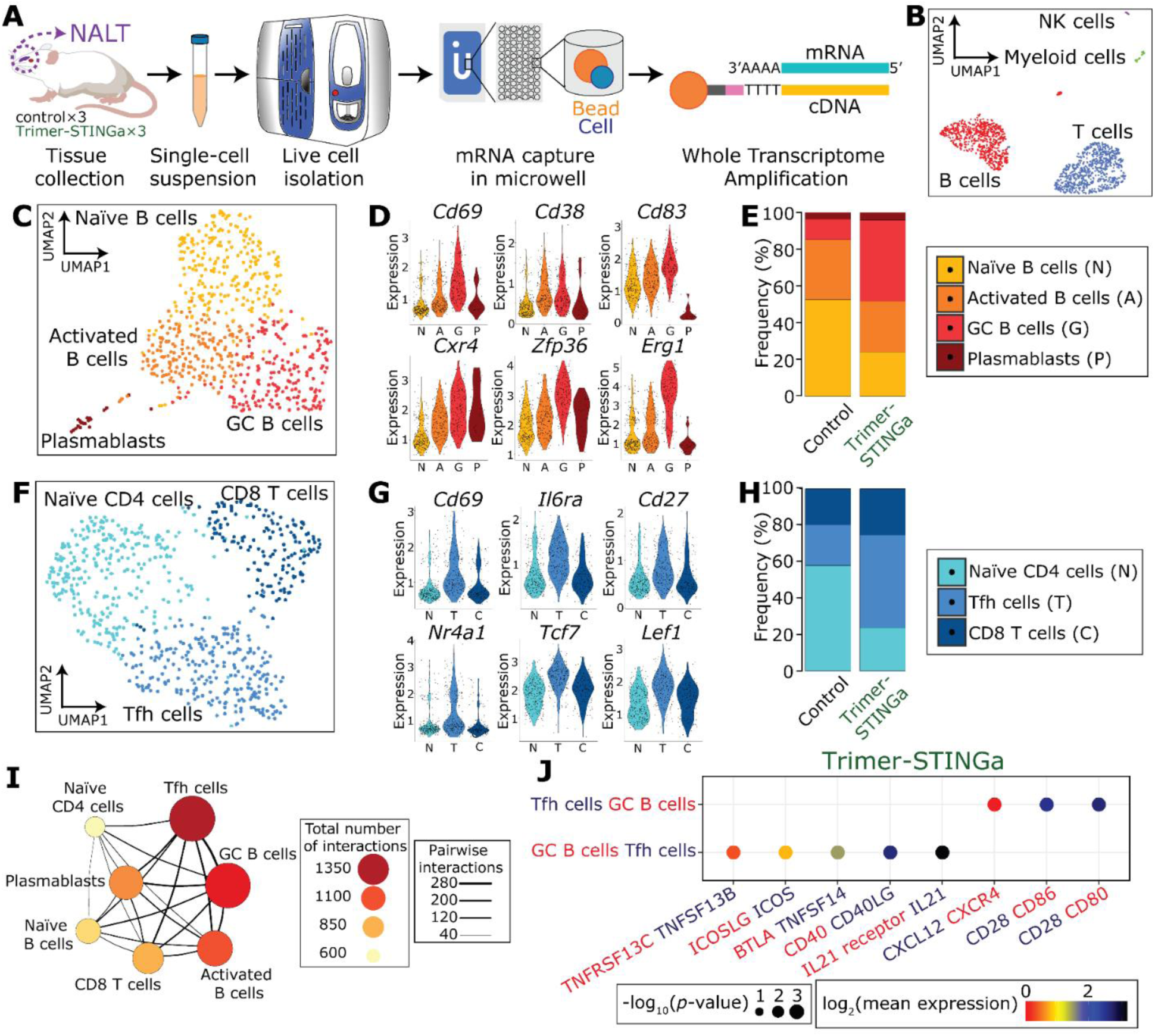
scRNA-seq confirms the nasal-associated lymphoid tissue (NALT) as an inductive site. A. Schematic of the experimental design for scRNA-seq on the NALT. B. Uniform manifold approximation and projection (UMAP) of the NALT immune cell profiles. C. Four clusters of B cells were identified based on UMAP: naïve, activated, and germinal center (GC) B cells; and plasmablasts. D. Violin plots of the relative expression of *Cd69, Cd38, Cd83, Cxcr4, Zfp36*, and *Erg1* in each of the four B-cell clusters. E. Bar plot illustrating the relative frequencies of each of the B-cell clusters in the NALT comparing the control and Trimer-STINGa groups. F. Three clusters of T cells were identified based on UMAP: naïve and follicular helper CD4 T cells; and CD8 T cells. G. Violin plots of the relative expression of *Cd69, Il6ra, Cd27, Nr4a1, Tcf7, and Lef1* in each of the three T-cell clusters. H. Bar plot illustrating the relative frequencies of the different T-cell clusters in the NALT comparing the control and Trimer-STINGa groups. I. Cell-cell interaction network illustrating the interactions between the immune cells in the NALT. The size and color of the circles reflect interaction counts, large and darker circles indicates more interaction with other cell types. The thickness of the connecting lines reflects the relative number of interactions between each pair of cells. J. Predicted ligand-receptor interactions between the GC B cells and the Tfh cells within the NALT. The relevant ligand-receptor pairs are shown as bubble plots.

We identified four B cell clusters: naïve B cells expressing *Cd19, Ms4a1 (CD20), Ighm*; germinal center B cells (GC B cells) expressing *Cd69, Cd38, Cd83, Cxcr4*, Z*fp36, Egr1, Erg3*, and *Irf4;* an intermediate B cell cluster comprising *of* activated B cells expressing *Cd38*; and ASC (plasmablasts) expressing *Sec61b, Casp3, and Tram1* but lacking expression of *Ms4a1 (CD20)* [Figure 3C-D and S6]^20,21^. Consistent with the role of the NALT as an inductive but not an effector site, we detected only a very small subpopulation of IgA (*Igha*) expressing cells, at least at this early time point (d15) [Figure S6]. Comparisons of the control and Trimer-STINGa groups showed a robust increase in the frequency of GC B cells with a concomitant decrease in naïve B cells [Figure 3E]. These results suggested that successful intranasal vaccination promoted activation of B cells similar to GCs, and we next investigated if T cells within the NALT supported B-cell differentiation. We identified three clusters within the T cells: one CD8^+^ T cell subpopulation expressing *Cd8a*, and two CD4^+^ T cell subpopulations (Figure 3F and S7). The CD4^+^ T cells were classified as naïve T cells (naïve) expressing *Cd4* and *Npm1;* and T follicular helper like (TFH) expressing *Cd69, Il6ra, Nr4a1 (Nur77), Tcf7* (TCF1), and *Lef1*, and also the memory markers *Cd27* and *Cd28* (Figure 3G and S7)^22-24^. The prominent difference in the control and the Trimer-STINGa groups was an increase in the ratio of Tfh/naïve CD4^+^ T cells (Figure 3H).

Since GC B cells and Tfh cells represented the dominant clusters in the Trimer-STINGa NALT, we investigated the nature of interactions between these cells in greater detail^25^. First, we visualized cell-cell interaction between the different B and T cell clusters in the NALT. We identified that Tfh cells were the dominant interacting cell type and interacted strongly with the GC B cell cluster (Figure 3I). At the molecular level, several well-documented receptor-ligand pairs, *Cd40*-*Cd40l, Il21r*-*Il21, Icosl*-*Icos*, and *Baffr* (*Tnfrsf13c*)-*Baff* (*Tnfsf13b*) were detected reciprocally on the GC B cells and the Tfh cells within the NALT (Figure 3J). These results showed that upon immunization with the Trimer-STINGa, the NALT promoted a GC-like T-cell dependent activation and differentiation of B cells which in turn can lead to long-lasting immunity.

### T-cell responses in the lung and IgA responses in the nasal compartment

We wanted to evaluate if the monomeric S protein could also elicit a comprehensive immune response. We used a monomeric version of the S protein containing mutations to the Furin binding site and a pair of stabilizing mutations (Lys986Pro and Val987Pro) [Figure 4A and S7]. SDS-PAGE of the monomeric protein showed a band between 130-180 kDa (Figure 4B). We immunized four mice with the monomeric S protein and the adjuvant (Monomer-STINGa) and again confirmed no weight loss in these animals (Figure S8). At d15, 100 % of the animals seroconverted, and the mean serum concentration of the anti-S IgG antibodies was 1:750 (Figure 4C). We confirmed that the serum anti-S antibodies were neutralizing with a mean ID50 of 1:188 [Figure 4D]. We evaluated T-cell responses with the same pool of immunodominant peptides. At d15, all four animals immunized with the Monomer-STINGa showed robust splenic T cell responses with a mean of 100 IFNγ spots/10^6^ cells (Figure 4E).

**Figure 4.**
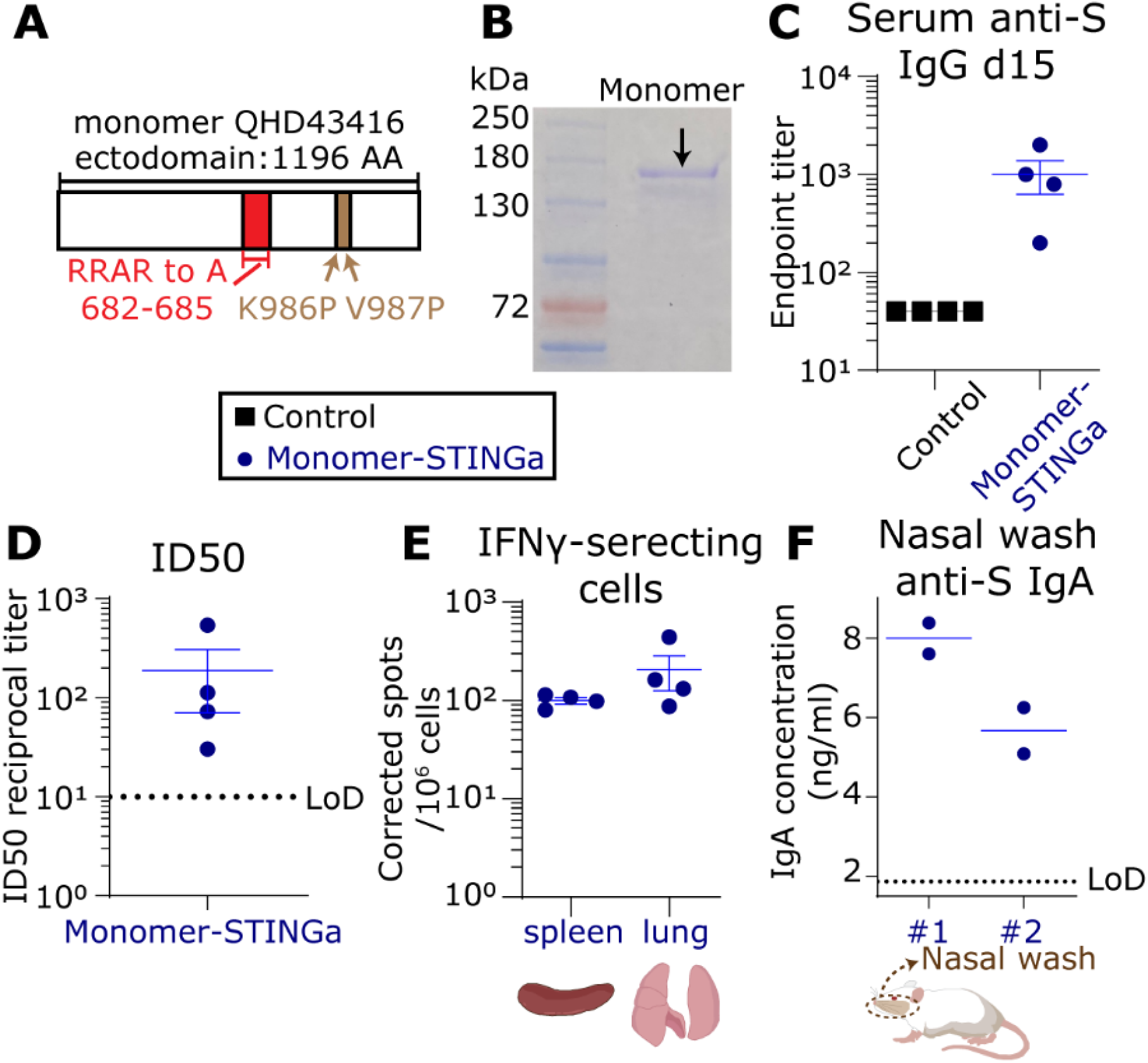
Systemic and mucosal responses elicited upon vaccination with Monomer-STINGa. A. Schematic of the monomeric protein used for immunization. B. Denaturing SDS-PAGE gel of the purified monomer protein. C. Humoral immune responses in the serum were evaluated using S-protein based IgG ELISA at day 15 after immunization. D. The ID50 of the serum antibody responses were measured using a pseudovirus neutralization assay E. Cellular immune responses in the spleen and lung were assessed using IFNγ ELISPOT assays. F. S-protein specific IgA was detectable in nasal wash of the animals in the Monomer-STINGa group For (C-F) the line represents the mean, and the error bars represent the standard error. LoD represents limit of detection of assay.

Animal models have shown that T cells in the lung are necesary for protection against pulmonary infection by respiratory pathogens^26^. Accordingly, we evaluated S-protein specific T-cell responses in the lung of the vaccinated animals. T-cell responses in the lung on d15 were detected at a mean of 206 IFNγ spots/10^6^ cells (Figure 4E). Collectively, these results established that intranasal administration using the Monomer-STINGa also elicited robust serum neutralizing antibodies and T-cell responses in both the spleen and the lung.

We finally investigated the antibody responses in the nasal compartment. We detected total IgA (ELISA) in the nasal wash from two animals. Both these nasal washes also had detectable anti-S IgA antibodies at a mean concentration of 7 ng/ml (Figure 4F). Consistent with the ability of the Monomer-STINGa to elicit mucosal immune responses, we also confirmed the detection of S-specific IgA secreting ASCs in the spleen of these mice (Figure S9). These results established that vaccination with the Monomer-STINGa elicits systemic immunity, T-cell responses in the lung and speen, and mucosal IgA responses.

## Discussion

Almost all the vaccine candidates that have advanced in the clinical trials of COVID19 are based on the intramuscular injection of DNA; nanoparticles loaded mRNA or viral vectors^3,27^. These methods have shown to elicit neutralizing antibodies in the serum of preclinical models and humans^27-29^. A fundamental limitation of all of these approaches is that they are not designed to elicit mucosal immunity. As prior work with other respiratory pathogens like influenza has shown, sterilizing immunity to virus re-infection requires adaptive immune responses in the respiratory tract and the lung^17,30,31^. In the context of COVID19 existing data supports that initial infection in the nasal compartment promotes/facilitates subsequent seeding of the virus to the lung^13^. The ability of vaccines to thus promote immunity at the mucosal sites and specifically within the nasal compartment can prevent seeding of the initial reservoir and control human transmission.

Intranasal vaccination is an attractive platform to elicit systemic and mucosal immunity. The fundamental challenge in intranasal vaccination is the ability to balance safety while ensuring immunogenicity leading to sterilizing immunity. Intranasal administration of live-attenuated vaccines in humans is hampered by concerns of safety^32^ and the use of the adenovirus vectored vaccines can be hampered by the presence of pre-existing immunity^33^. Subunit vaccines are attractive candidates that do not suffer from these drawbacks. However, the ability of subunit vaccines to elicit potent and sterilizing immune responses is critically dependent on the choice of appropriate adjuvants.

We demonstrate that STINGa encapsulated in liposomes can function as potent adjuvants. A single intranasal immunization with protein S and the adjuvant can elicit neutralizing antibodies in the serum comparable to other vaccine candidates. We established that intranasal vaccination leads to IgA responses in the lung and directly in the nasal compartment, and we detected B cells secreting IgA in the spleen. We show using scRNA-seq that the NALT functions as an inductive site upon intranasal vaccination, leading to co-ordinated activation/differentiation of B and T cells resembling germinal centers. Moreover, intranasal vaccination also induced S-specific T cell responses in both the spleen and locally in the lung. Collectively, these results illustrate the advantages of optimal intranasal immunization. While we demonstrate cellular and humoral immunity both systemically and locally in the respiratory tract, future studies using appropriate challenge models in non-human primates are required to establish whether we elicit sterilizing immunity and to test the durability of our responses longitudinally. However, we note that due to the severity of the pandemic, most vaccines currently in clinical trials have advanced without testing in preclinical challenge models. Although we have shown the immunity elicited upon a single-dose, we can add a booster dose to increase the durability of the responses, if appropriate.

In summary, our study establishes that intranasal spike subunit vaccines with liposomal STINGa as adjuvants is safe and elicits comprehensive immunity against SARS-CoV-2. In the context of a pandemic, the intranasal vaccine has two compelling advantages. First, the easy access to the nasal cavity makes intranasal administration non-invasive. It is particularly suited for mass vaccinations of large cohorts, including elderly patients and children with minimal clinical infrastructure. Second, from the standpoint of disease control, the ability to control SARS-CoV-2 infection at the first point of entry in the nasal compartment and before spreading to the lung is a desirable option to halt disease progression in individuals and disease transmission across populations. We suggest that our promising vaccine candidate supports human testing.

## Methods

### Lipids and STINGa

We purchased the STINGa, 2′-3’′cyclic guanosine monophosphate adenosine monophosphate (cGAMP) from Chemietek (Indianapolis, IN). 1,2-dipalmitoyl-sn-glycero-3-phosphocholine (DPPC), 1,2-dipalmitoyl-sn-glycero-3-phospho-(1’-rac-glycerol) (DPPG), and 1,2-dipalmitoyl-sn-glycero-3-phosphoethanolamine-N-[methoxy(polyethylene glycol)-2000] (DPPE-PEG2000) were obtained from Avanti Polar Lipids (Alabaster, AL). Cholesterol was obtained from Sigma Aldrich (St. Louis, MO).

### Preparation of STINGa loaded liposomes and vaccine formulation

The liposomes were composed of a molar ratio of 10:1:1:1 of DPPC, DPPG, Cholesterol (Chol), and DPPE-PEG2000. To prepare the liposomes, we mixed the lipids in CHCl_3_ and CH_3_OH, and the solution was evaporated by a vacuum rotary evaporator for approximately 80 min at 45 °C. We dried the resulting lipid thin film until all organic solvent was evaporated. We hydrated the lipid film by adding a pre-warmed cGAMP solution (0.3 mg/ml in PBS buffer at pH 7.4). The hydrated lipids were mixed at elevated temperature 65 °C for an additional 30 min, then subjected to freeze-thaw cycles. We sonicated the mixture for 60 min using a Brandson Sonicator (40 kHz). The free untrapped cGAMP was removed by Amicon Ultrafiltration units (MW cut off 10kDa). We washed the cGAMP-liposomes three times using PBS buffer. The cGAMP concentration in the filtrates was measured by Take3 Micro-Volume absorbance analyzer of Cytation 5 (BioTek) against a calibration curve of cGAMP at 260 nm. We calculated the final concentration of liposomal encapsulated cGAMP and encapsulation efficiency by subtracting the concentration of free drug in the filtrate.

We mixed the S protein monomer (4 µg) or the trimer (10 µg) with the STINGa-liposomal suspensions at room temperature to allow the adsorption of the protein onto the liposomes. The formulated vaccine was stored at 4 °C and used for up to 2 months. The average particle diameter, polydispersity index, and zeta potential were characterized by Litesizer 500 (Anton Paar) at room temperature.

### Mice and immunization

All the animal experiments were reviewed and approved by UH IACUC. Female, 7-9 week-old BALB/c mice were purchased from Charles River Laboratories. Prior to immunization, we anesthetized the mice by intraperitoneal injection of ketamine and xylazine. We immunized the mice intranasally with different formulations: (1) the adjuvant only group was administrated with liposome-STINGa; (2) the control group was administrated with protein only; (3) the Trimer-STINGa group was administrated with protein and liposome-STINGa; and (4) the Monomer-STINGa group was administrated with protein and liposome-STINGa. The monomeric and trimeric protein was obtained from BEI Resources (VA, USA) and Creative BioMart (NY, USA), respectively.

### Body weight monitoring and sample collection

The body weight of the animals was monitored every 2-3 days over two weeks after immunization. Sera were collected seven days and 15 days after post-vaccination for detection of the humoral response. Nasal wash, BALF, NALT, lung, and spleen were harvested and processed 15 days after the administration, essentially as previously described^34,35^. Sera and other biological fluids (with protease inhibitors) were kept at -80 °C for long-term storage. After dissociation, the splenocytes and lung cells were frozen in FBS+10% DMSO and stored in the liquid nitrogen vapor phase until further use.

### ELISA

Anti-S protein antibody titers in serum or other biological fluids were determined using ELISA. Briefly, we coated 1 μg/ml spike protein (Sino Biological, PA, USA) onto ELISA plates (Corning, NY, USA) in PBS overnight at 4 °C or 2 hours at 37 °C. The plate was blocked with PBS+1%BSA (Fisher Scientific, PA, USA)+0.1%Tween20™ (Sigma-Aldrich, MD, USA) for 2 hours at room temperature. After washing, we added the samples at different dilutions. We detected the captured antibodies by HRP-conjugated anti-mouse IgG (Jackson ImmunoResearch Laboratories, 1 in 5,000; PA, USA), anti-mouse IgA (Bethyl Laboratories, 1 in 10,000; TX, USA) and detection antibody against mouse IgA (1 in 250) from the mouse total IgA ELISA kit from Invitrogen (CA, USA). The positive control (anti-S IgG) was obtained from Abeomics (CA, USA).

### ELISPOT

IFNγ ELISpot assay was performed using Mouse IFN-γ ELISPOT Basic kit (ALP), following the manufacture’s instructions (Mabtech, VA, USA). Frozen splenocytes or lung cells were thawed and seeded on the ELISPOT plate (100-300 k cells per well) without further culturing. The splenocytes were incubated with the spike protein peptide pool at 1.5 μg/ml/peptide (Miltenyi Biotec, Germany) at 37 °C for 16-18 hours. The ELISPOT plates were read using a ImmunoSpot® S6 MICRO analyzer.

IgA-secreting cells were detected using Mouse IgG/IgA Double-Color ELISPOT from Immunospot (OH, USA) following the manufacture’s instruction. For the total IgA-secreting cells in the spleen, the splenocytes were thawed and seeded to the capture antibody coated ELISPOT plate immediately. The cells were incubated at 37 °C for 16-18 hours, followed by the development. For the anti-S IgA producing cells, thawed splenocytes were cultured in complete media [RPMI-1640 (Corning, NY, USA) +10% FBS (R&D System, MN, USA), 100 μg/ml NormocinTM (InvivoGen, CA, USA), 2 mM L-Glutamine, 1 mM sodium pyruvate, 10 mM HEPES] and B-Poly-STM (1:1000 dilution) at 4 million cells/ml. The wells were coated with 5 μg/ml of the spike protein (Sino Biological, PA, USA) overnight at 4 °C. The spleen cells were washed and seeded onto the plate at 37 °C for 16-18 hours.

### Cell lines and plasmids

293T cells stably expressing SARS-CoV-2 receptor human angiotensin-converting enzyme II (ACE2) and plasma membrane-associated type II transmembrane serine protease, TMPRSS2 (293 T/ACE2-TMPRSS2) were a generous gift from Dr. Siyan Ding (Washington University School of Medicine, St. Louis, MO, USA)^36^. The cells were cultured in Dulbecco’s modified Eagle’s medium (DMEM) supplemented with 10% fetal bovine serum. The expression plasmids for SARS-CoV-2 S protein pCAGGS Containing the SARS-CoV-2, Wuhan-Hu-1 Spike Glycoprotein Gene was obtained from BEI Resources (Manassas, VA). Plasmids encoding GFP expressing Lentiviral vector, helper plasmids pMDLg/pRRE,pRSV-Rev, and VSV-G protein-encoding plasmid pMD2.G were obtained from Addgene (Watertown, MA).

### Generation of SARS-CoV-2 spike protein-expressing reporter virus

To determine the titer of neutralizing antibodies in the serum of immunized mice, SARS-CoV-2-S pseudotyped lentiviral system was used as a surrogate for SARS-CoV-2 infection^8^. Pseudotyped viral (PsV) stocks were generated by co-transfecting stable ACE-2 and TMPRSS2 expressing 293 T cells with pLVX-AcGFP1-C1, pMDLg/pRRE, pRSV-Rev and viral envelope protein expression plasmids pCAGGS containing the SARS-CoV-2, Wuhan-Hu-1 Spike Glycoprotein Gene or VSV-G envelope expressing plasmid pMD2.G using the approach employed by Wu *et al*.^*8*^ and Almasaud *et al*. ^37^ to generate the PsV particles. The PsV particles in supernatants were harvested 48 h post-transfection, filtered and stored at −80°C as described previously^37,38^. The dose titer of PsV was determined by infecting ACE-2 and TMPRSS2 expressing 293T cells for 48 h and using Celigo imaging system for imaging and counting virus infected fluorescent cells^39^. The viral titers were expressed as fluorescent focus forming units (FFU)/well^40,41^.

### Neutralizing Antibody (Nab) Titration Assay for SARS-CoV-2

For microneutralization assay, ACE2-TMPRSS2 expressing 293 T cells were cultured overnight in a half area 96-well plate compatible with Nexcelom Celigo imager at a concentration of 1 ×10^4^ cells per well in 100 µl of complete media. On the day of the assay, heat-inactivated serum from mice was thawed and diluted 1:20 to 1:640 in a six point, two-fold series in serum-free DMEM. In a 96 well plate, 50 µl of diluted serum was mixed with 50 µl of GFP expressing SARS-CoV-2 spike expressing PsV (150-300 FFU/well) and incubated at 37 °C for 1 hour. After 1 h, this mixture of added to ACE2-TMPRSS2 expressing 293 T cells and incubated for 48 h. The infection of target cells was determined by imaging and counting FFU/well using Celigo imaging system. Each sample was tested in triplicate wells. SARS-CoV-2 Spike S1 rabbit Mab (Clone #007, Sino Biological, Wayne, PA) was used as a positive control for neutralizing activity and VSV-G expressing PsV was used as a negative control for the specificity of neutralization function. The 50% inhibitory dose (ID50) titers of NAbs were calculated using nonlinear regression (GraphPad Prism).

### NALT Collection and Fluorescence-Activated Cell Sorting

We isolated the NALT from the mice after euthanasia precisely as described previously^34,42^. We lysed the red blood cells by incubating the cells in ACK lysis buffer (Thermo Fisher Scientific, Waltham, MA) for 3 minutes. We subsequently washed the single-cell suspensions with PBS, resuspended them in PBS + 2% FBS, and added 50 nM Helix NP Blue (BioLegend, San Diego, CA) detection of live/dead cells. We used a BD FACSMelody cell sorter (BD Biosciences, San Jose, CA) to collect live cells.

### Single-Cell Library Preparation and Sequencing

We labeled each group of NALT cells separately with the Sample-Tags from the BD Mouse Immune Single-Cell Multiplexing Kit (BD Biosciences, San Jose, CA), described in “Single Cell Labeling with the BD Single-Cell Multiplexing Kits” protocol. We then proceeded to library preparation with a mixture of ∼6000 cells (3000 cells from each group). We prepared the whole transcriptome using the BD Rhapsody System following the BD “mRNA Whole Transcriptome Analysis (WTA) and Sample Tag Library Preparation Protocol”. We assessed the quality and quantity of the final library by Agilent 4200 TapeStation system using the Agilent High Sensitivity D5000 ScreenTape (Agilent Technologies, Santa Clara, CA) and a Qubit Fluorometer using the Qubit dsDNA HS Assay, respectively. We diluted the final library to 3 nM concentration and used a HiSeq PE150 sequencer (Illumina, San Diego, CA) to perform the sequencing.

### Quantification and statistical analysis

#### Analyzing WTA data via Seven Bridges

We uploaded and analyzed the FASTQ files on Seven Bridges website (Seven Bridges Genomics) by running the “BD Rhapsody WTA Analysis Pipeline” (BD Biosciences, San Jose, CA). After performing alignment, filtering, and sample tag detection, we downloaded and used the pipeline’s final outputs, including the sample tag calls and molecule count information for further analysis in R (version 4.0.1) using Seurat Package (version 3.0)^43^.

#### Downstream analysis of WTA data in R

We first used the SAVER Package to recover the gene expression and provide a reliable quantification of low expressed genes across the data^44^. By following the standard processing workflow in Seurat Package, we acquired the clustering and gene expression data. We removed cells with < 1000 Unique Molecular Identifiers (UMIs) and high mitochondrial gene expression (> 20% of the reads), and we ended up with 1398 single-cell profiles (660 control, 738 treated) with a mean UMI of 1648.

To analyze the cell-cell communication at the molecular level, we used the recovered molecule counts by SAVER in CellPhoneDB analysis tool^25^. First, we transformed the mouse genes to their human orthologous using BiomaRt Package (version 2.38.0)^45^. Then, we categorized all T cells and B cells by their subpopulations and group (control and Trimer-STINGa) into 14 cell types. According to statistical tests calculated CellPhoneDB, we filtered out the ligand-receptor pairs with p values > 0.05 and evaluated the relationship between different cell types with the significant pairs. To generate the network of interactions, we applied igraph Package (version 1.2.5)^46^.

## Acknowledgments

This publication was supported by the NIH (U01AI148118) and Owens foundation. XL acknowledges partial funding support from the National Cancer Institute (NIH R15CA182769, P20CA221731, P20CA221696) and CPRIT (RP150656). The following reagent was produced under HHSN272201400008C and obtained through BEI Resources, NIAID, NIH: Spike Glycoprotein (Stabilized) from SARS-Related Coronavirus 2, Wuhan-Hu-1, Recombinant from Baculovirus, NR-52308. Supported by the NIH/NCI under award number P30 CA016672 and used the M.D. Anderson ORION core. We would like to acknowledge Prof. Shaun Zhang for sharing guidance on animal protocols; Drs. Ankita Leekha and Irfan Bandey for assistance with animal experiments; and Prof. Cliona Rooney for sharing ELISPOT plates. We thank BD for the generous loaner of a FACS Melody and Rhapsody; and Intel for the generous loaner of a cluster.

## Author contributions

NV conceived the study. XA, MM, XL, and NV designed the study. XA, MM, AR, MF, SB, SS, and XL performed experiments. XA, SS, XL, and SB analyzed the data. AR and MF performed bioinformatic analyses. XA, SS, XL, MF, AR, CY, and NV made figures and wrote the manuscript.

## Supplementary Figures

**Figure S1.**
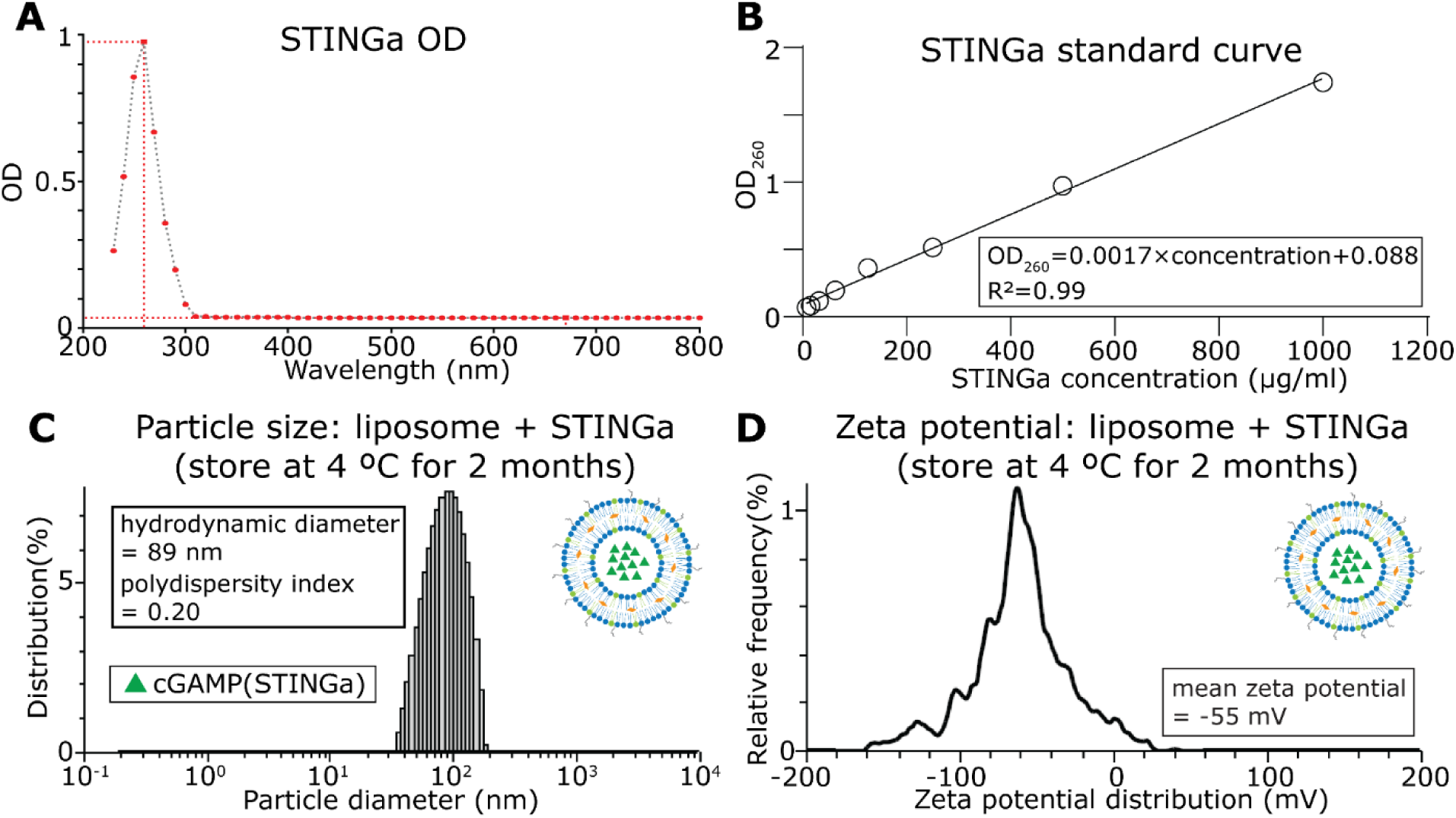
Quantification of STINGa and characterization of the stability of STINGa-liposomes. A. UV-visible absorption spectrum of cGAMP (STINGa) showing the absorption maximum at 260 nm. B. The standard curve used for calculating the concentration of free STINGa. C. Distribution of liposomal particle sizes measured by dynamic light scattering (DLS) after storage at 4 °C for two months. D. Zeta potential of the liposomes measured by electrophoretic light scattering (ELS) after storage 4 °C for two months.

**Figure S2.**
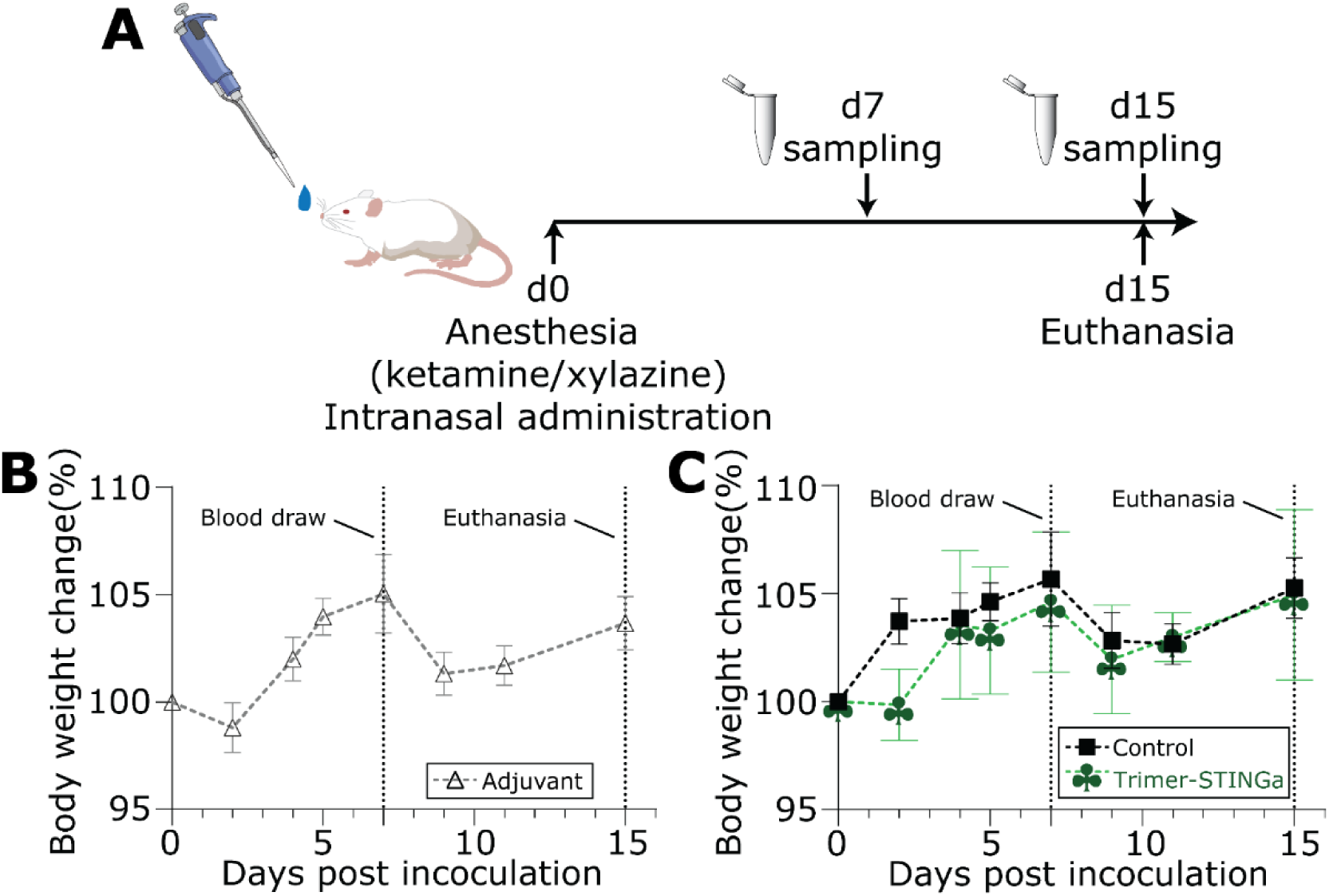
Immunization and monitoring of mice. A. Overall timeline for the immunization and sample collection B-C. Monitoring of the body weight of the mice administered: (B) liposomal-STINGa, and (C) either protein only or Trimer-STINGa

**Figure S3.**
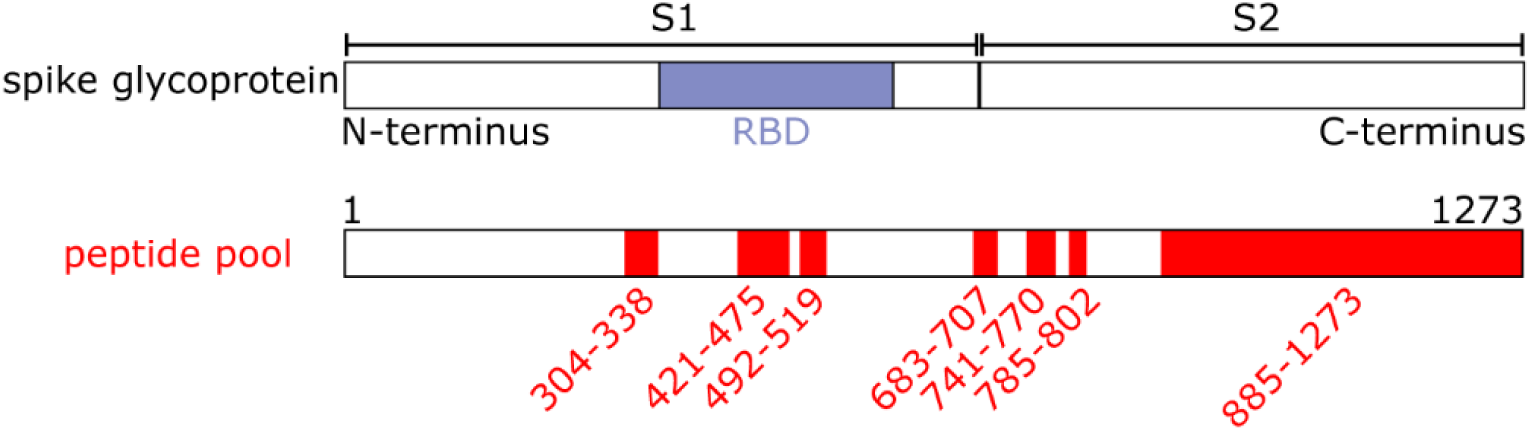
Protein level mapping of the conserved peptides (15mers) used for IFN-γ ELISPOT assays.

**Figure S4.**
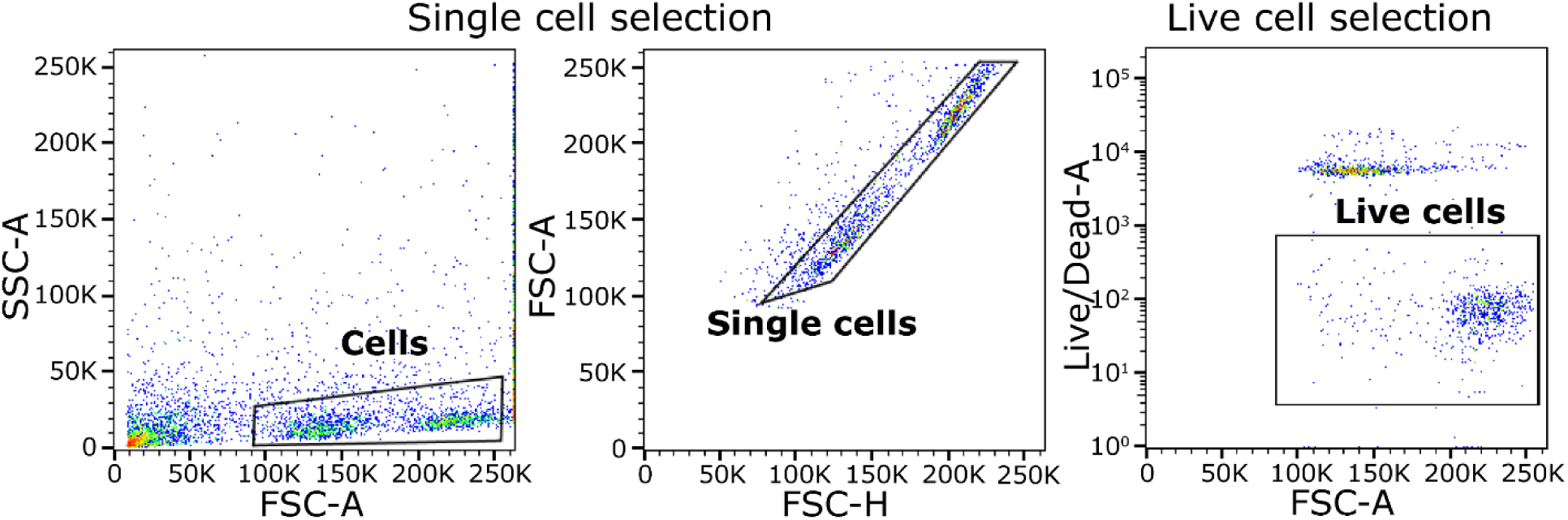
Flow cytometric gates used for the isolation of live cells for scRNA-seq.

**Figure S5.**
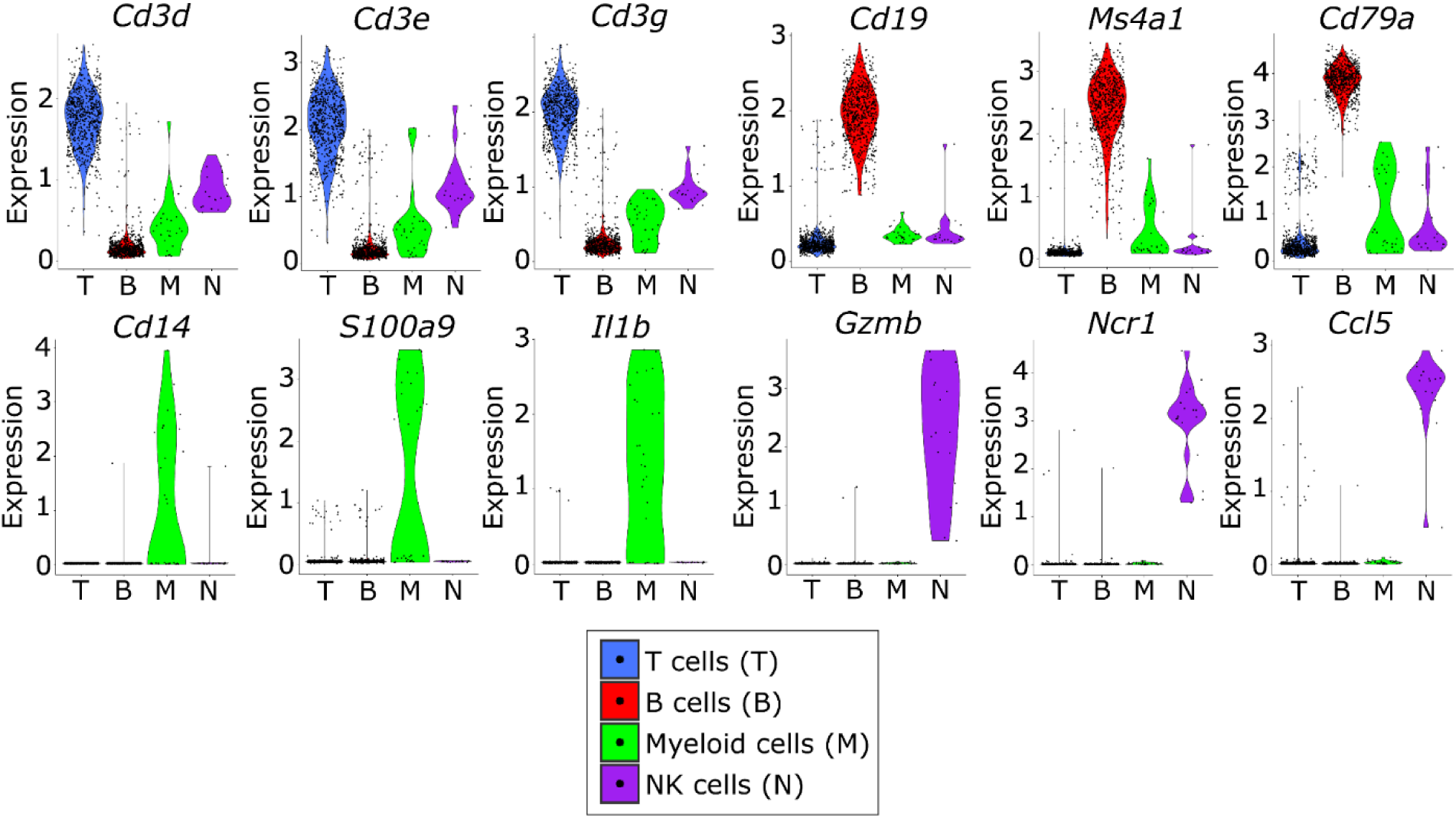
Violin plots of the relative expression of T-cell specific (*Cd3d, Cd3e*, and *Cd3g*), B-cell specific (*Cd19, Cd20*, and *Cd79*), myeloid specific (*Cd14, S100a9*, and *Il1b*), and NK-cell specific (*GzmB, Ncr1*, and *Ccl5*) transcripts.

**Figure S6.**
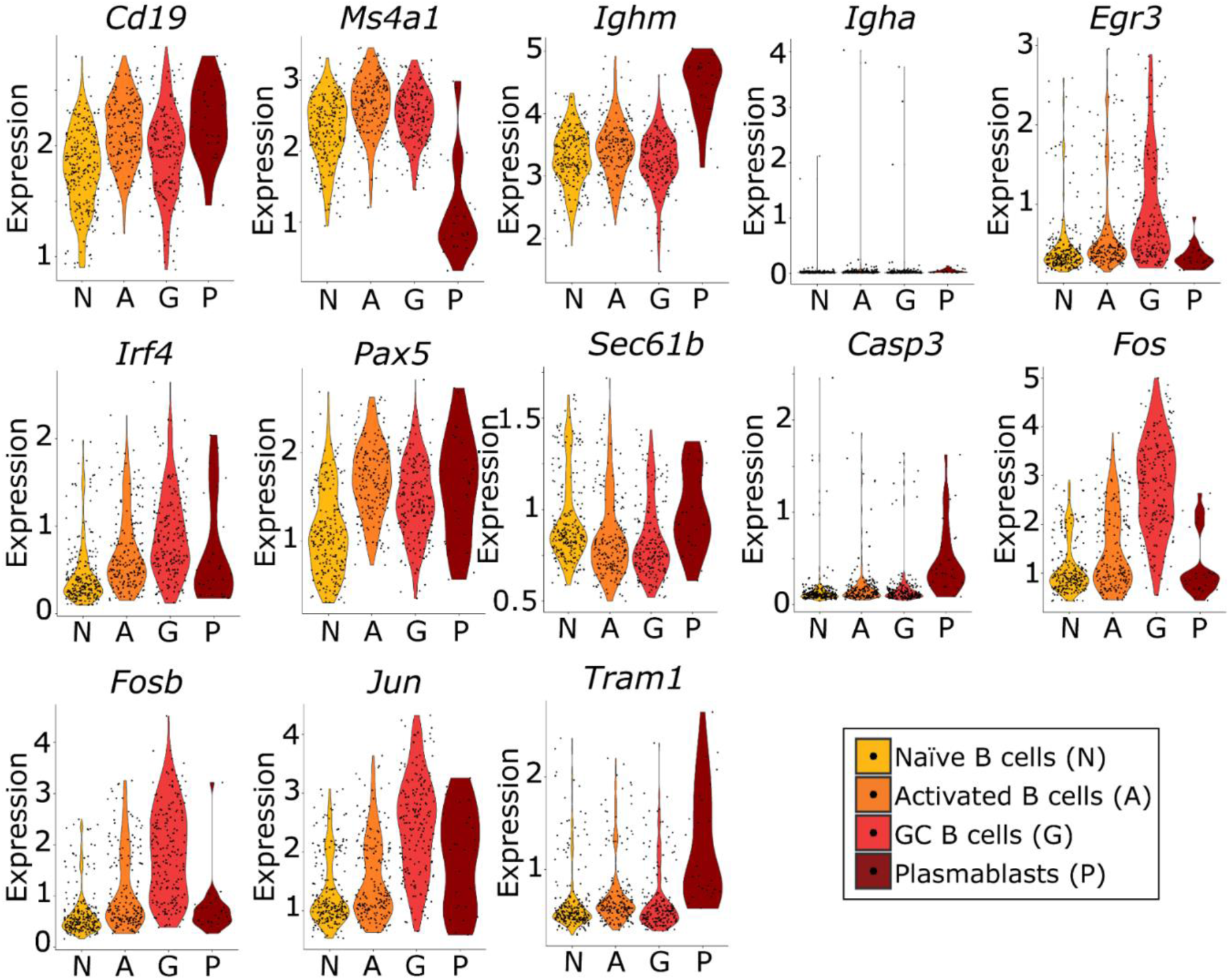
Violin plots of the relative expression of the key molecules associated with each of the four B-cell clusters.

**Figure S7.**
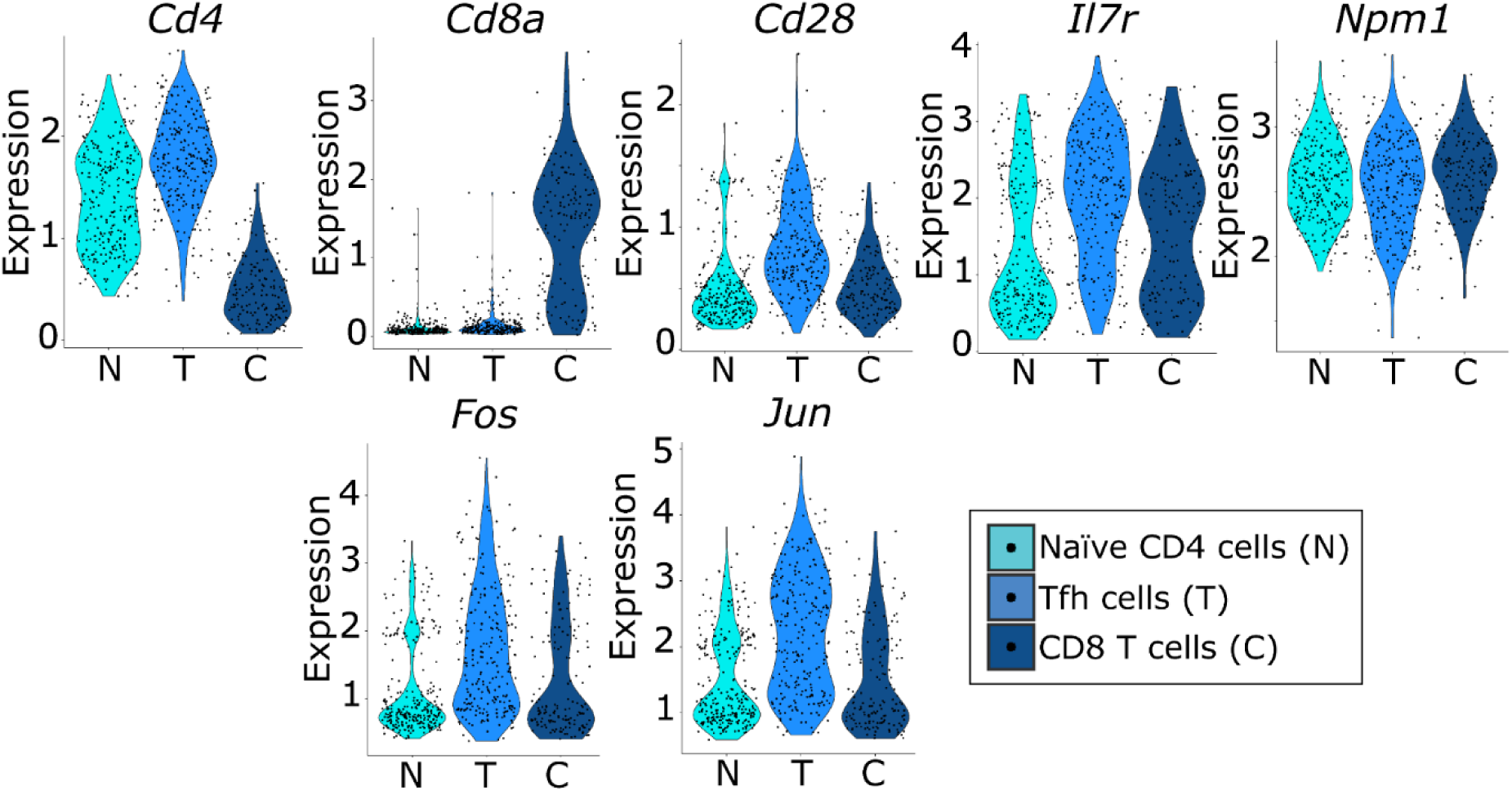
Violin plots of the relative expression of the key molecules associated with each the three T-cell clusters.

**Figure S8.**
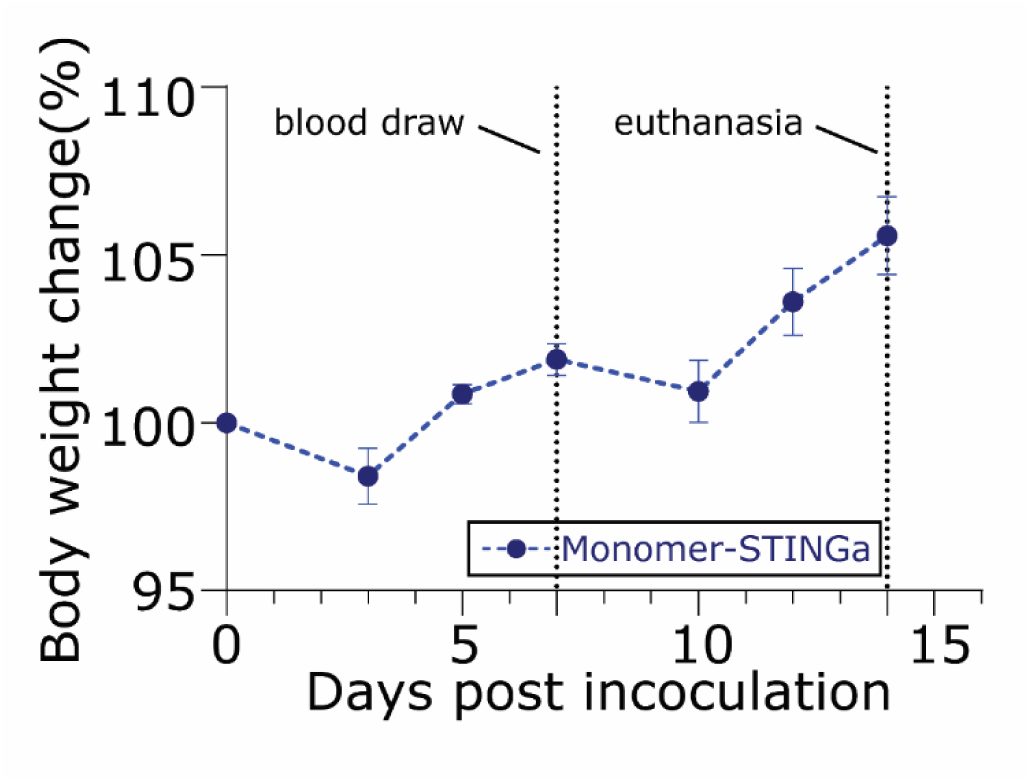
The Monomer-STINGa or the control protein does not adversely affect the mice after immunization. The circles denote the mean and error bars denote standard error.

**Figure S9.**
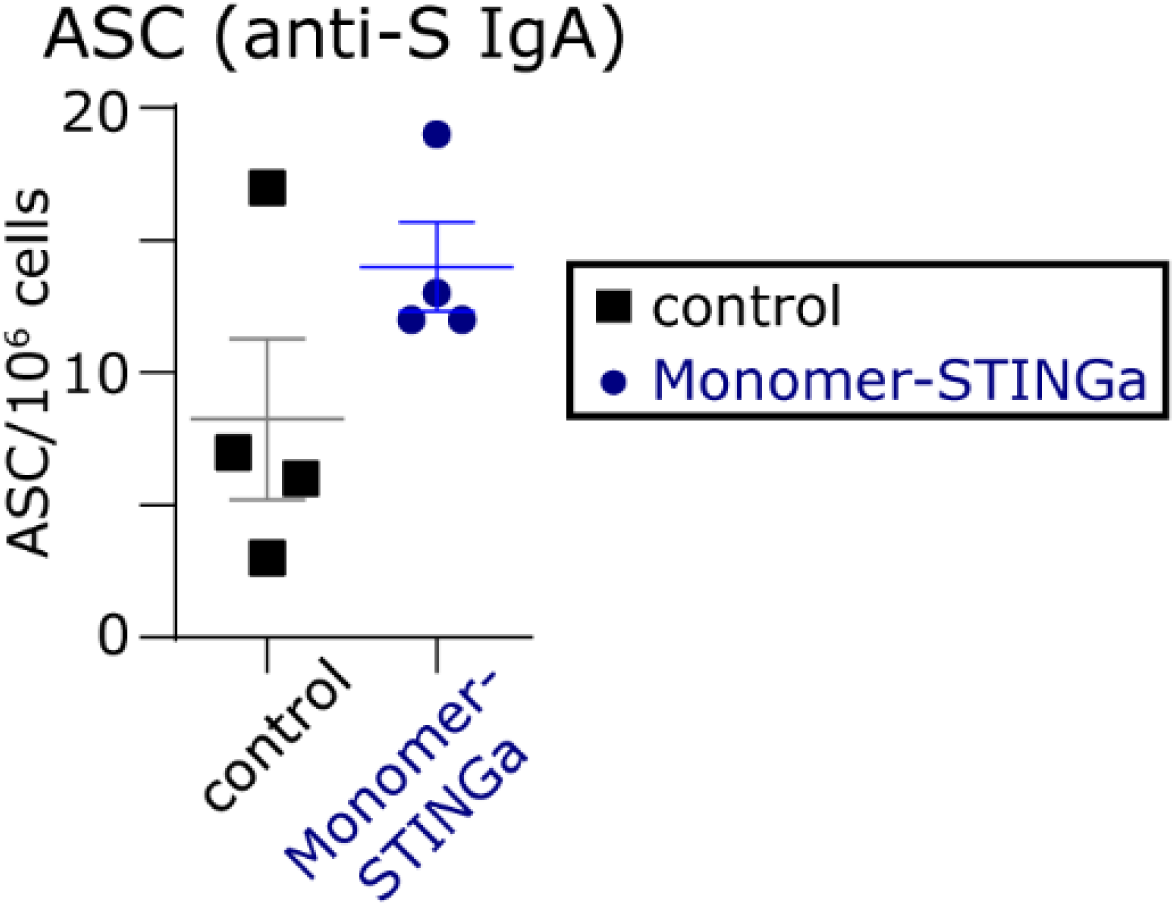
Antibody secreting cells (ASCs) secreting S-protein specific IgA in the spleen were detected using ELISPOT assays. The line denotes the mean, and the error bars denote the standard error.

## References

1 Hoffmann, M. et al. SARS-CoV-2 Cell Entry Depends on ACE2 and TMPRSS2 and Is Blocked by a Clinically Proven Protease Inhibitor. Cell 181, 271–280 e278, (2020).

2 Robbiani, D. F. et al. Convergent antibody responses to SARS-CoV-2 in convalescent individuals. Nature, (2020).

3 Smith, T. R. F. et al. Immunogenicity of a DNA vaccine candidate for COVID-19. Nat Commun 11, 2601, (2020).

4 Erasmus, J. H. et al. Single-dose replicating RNA vaccine induces neutralizing antibodies against SARS-CoV-2 in nonhuman primates. bioRxiv, (2020).

5 Long, Q. X. et al. Clinical and immunological assessment of asymptomatic SARS-CoV-2 infections. Nat Med, (2020).

6 Pollan, M. et al. Prevalence of SARS-CoV-2 in Spain (ENE-COVID): a nationwide, population-based seroepidemiological study. Lancet, (2020).

7 Ibarrondo, F. J. et al. Rapid Decay of Anti–SARS-CoV-2 Antibodies in Persons with Mild Covid-19. New England Journal of Medicine, (2020).

8 Wu, F. et al. Neutralizing antibody responses to SARS-CoV-2 in a COVID-19 recovered patient cohort and their implications. medRxiv, (2020).

9 Gallais, F. et al. Intrafamilial Exposure to SARS-CoV-2 Induces Cellular Immune Response without Seroconversion. bioRxiv, (2020).

10 Grifoni, A. et al. Targets of T Cell Responses to SARS-CoV-2 Coronavirus in Humans with COVID-19 Disease and Unexposed Individuals. Cell 181, 1489–1501 e1415, (2020).

11 Weiskopf, D. et al. Phenotype and kinetics of SARS-CoV-2-specific T cells in COVID-19 patients with acute respiratory distress syndrome. Sci Immunol5, (2020).

12 Braun, J. et al. Presence of SARS-CoV-2 reactive T cells in COVID-19 patients and healthy donors. bioRxiv, (2020).

13 Hou, Y. J. et al. SARS-CoV-2 Reverse Genetics Reveals a Variable Infection Gradient in the Respiratory Tract. Cell, (2020).

14 Dubensky, T. W., Jr., Kanne, D. B. & Leong, M. L. Rationale, progress and development of vaccines utilizing STING-activating cyclic dinucleotide adjuvants. Ther Adv Vaccines 1, 131–143, (2013).

15 Martin, T. L. et al. Sublingual targeting of STING with 3’3’-cGAMP promotes systemic and mucosal immunity against anthrax toxins. Vaccine 35, 2511–2519, (2017).

16 Luo, J. et al. Enhancing Immune Response and Heterosubtypic Protection Ability of Inactivated H7N9 Vaccine by Using STING Agonist as a Mucosal Adjuvant. Front Immunol 10, 2274, (2019).

17 Wang, J. et al. Pulmonary surfactant-biomimetic nanoparticles potentiate heterosubtypic influenza immunity. Science 367, (2020).

18 Anft, M. et al. COVID-19 progression is potentially driven by T cell immunopathogenesis. medRxiv, (2020).

19 Ahmed, S. F., Quadeer, A. A. & McKay, M. R. Preliminary Identification of Potential Vaccine Targets for the COVID-19 Coronavirus (SARS-CoV-2) Based on SARS-CoV Immunological Studies. Viruses 12, (2020).

20 Ellebedy, A. H. et al. Defining antigen-specific plasmablast and memory B cell subsets in human blood after viral infection or vaccination. Nat Immunol17, 1226–1234, (2016).

21 Ise, W. et al. T Follicular Helper Cell-Germinal Center B Cell Interaction Strength Regulates Entry into Plasma Cell or Recycling Germinal Center Cell Fate. Immunity 48, 702–715 e704, (2018).

22 Choi, Y. S. et al. LEF-1 and TCF-1 orchestrate T(FH) differentiation by regulating differentiation circuits upstream of the transcriptional repressor Bcl6. Nat Immunol16, 980–990, (2015).

23 Wu, H. et al. Molecular Control of Follicular Helper T cell Development and Differentiation. Front Immunol9, 2470, (2018).

24 Cano-Gamez, E. et al. Single-cell transcriptomics identifies an effectorness gradient shaping the response of CD4(+) T cells to cytokines. Nat Commun 11, 1801, (2020).

25 Vento-Tormo, R. et al. Single-cell reconstruction of the early maternal-fetal interface in humans. Nature 563, 347–353, (2018).

26 Pizzolla, A. et al. Influenza-specific lung-resident memory T cells are proliferative and polyfunctional and maintain diverse TCR profiles. J Clin Invest 128, 721–733, (2018).

27 van Doremalen, N. et al. ChAdOx1 nCoV-19 vaccination prevents SARS-CoV-2 pneumonia in rhesus macaques. bioRxiv, (2020).

28 Jackson, L. A. et al. An mRNA Vaccine against SARS-CoV-2 — Preliminary Report. New England Journal of Medicine, (2020).

29 Mulligan, M. J. et al. Phase 1/2 Study to Describe the Safety and Immunogenicity of a COVID-19 RNA Vaccine Candidate (BNT162b1) in Adults 18 to 55 Years of Age: Interim Report. bioRxiv, (2020).

30 Dutta, A. et al. Sterilizing immunity to influenza virus infection requires local antigen-specific T cell response in the lungs. Sci Rep 6, 32973, (2016).

31 Laurie, K. L. et al. Multiple infections with seasonal influenza A virus induce cross-protective immunity against A(H1N1) pandemic influenza virus in a ferret model. J Infect Dis 202, 1011–1020, (2010).

32 Zaman, M., Chandrudu, S. & Toth, I. Strategies for intranasal delivery of vaccines. Drug Deliv Transl Res 3, 100–109, (2013).

33 Fausther-Bovendo, H. & Kobinger, G. P. Pre-existing immunity against Ad vectors: humoral, cellular, and innate response, what’s important? Hum Vaccin Immunother 10, 2875–2884, (2014).

34 Cisney, E. D., Fernandez, S., Hall, S. I., Krietz, G. A. & Ulrich, R. G. Examining the Role of Nasopharyngeal-associated Lymphoreticular Tissue (NALT) in Mouse Responses to Vaccines. Journal of Visualized Experiments, (2012).

35 Van Hoecke, L., Job, E. R., Saelens, X. & Roose, K. Bronchoalveolar Lavage of Murine Lungs to Analyze Inflammatory Cell Infiltration. Journal of Visualized Experiments, (2017).

36 Zang, R. et al. TMPRSS2 and TMPRSS4 promote SARS-CoV-2 infection of human small intestinal enterocytes. Sci Immunol5, (2020).

37 Almasaud, A., Alharbi, N. K. & Hashem, A. M. Generation of MERS-CoV Pseudotyped Viral Particles for the Evaluation of Neutralizing Antibodies in Mammalian Sera. Methods Mol Biol 2099, 117–126, (2020).

38 Wu, F. et al. Neutralizing antibody responses to SARS-CoV-2 in a COVID-19 recovered patient cohort and their implications. medRxiv, 2020.2003.2030.20047365, (2020).

39 Masci, A. L. et al. Integration of Fluorescence Detection and Image-Based Automated Counting Increases Speed, Sensitivity, and Robustness of Plaque Assays. Mol Ther Methods Clin Dev 14, 270–274, (2019).

40 Baker, S. F., Nogales, A., Santiago, F. W., Topham, D. J. & Martinez-Sobrido, L. Competitive detection of influenza neutralizing antibodies using a novel bivalent fluorescence-based microneutralization assay (BiFMA). Vaccine 33, 3562–3570, (2015).

41 Shambaugh, C. et al. Development of a High-Throughput Respiratory Syncytial Virus Fluorescent Focus-Based Microneutralization Assay. Clin Vaccine Immunol24, (2017).

42 Asanuma, H. et al. Isolation and characterization of mouse nasal-associated lymphoid tissue. J Immunol Methods 202, 123–131, (1997).

43 Stuart, T. et al. Comprehensive Integration of Single-Cell Data. Cell 177, 1888–1902 e1821, (2019).

44 Huang, M. et al. SAVER: gene expression recovery for single-cell RNA sequencing. Nat Methods 15, 539–542, (2018).

45 Durinck, S. et al. BioMart and Bioconductor: a powerful link between biological databases and microarray data analysis. Bioinformatics 21, 3439–3440, (2005).

46 Csardi, G. & Nepusz, T. The igraph software package for complex network research. InterJournal Complex Systems, 1695, (2006).

